# Cancer Associated Fibroblasts Mediate Cancer Progression and Remodel the Tumouroid Stroma

**DOI:** 10.1101/859041

**Authors:** Judith Pape, Tarig Magdeldin, Katerina Stamati, Agata Nyga, Marilena Loizidou, Mark Emberton, Umber Cheema

## Abstract

**Objective:** Cancer associated fibroblasts (CAFs) are highly differentiated and heterogenous cancer stromal cells that promote tumour growth, angiogenesis and matrix remodelling.

**Design:** We utilised a novel 3D in vitro model of colorectal cancer, composed of a cancer mass and surrounding stromal compartment. We compared cancer invasion with an acellular stromal surround, a ‘healthy’ or normal cellular stroma and a cancerous stroma. For the cancerous stroma we incorporated six patient-derived CAF samples to study their differential effects on cancer growth, vascular network formation, and remodelling.

**Results:** CAFs enhanced the distance and surface area of the invasive cancer mass whilst inhibiting vascular-like network formation. These processes were driven by the upregulation of hepatocyte growth factor (HFG), metallopeptidase inhibitor 1 (TIMP1) and fibulin 5 (FBLN5).

Remodelling appeared to occur through the process of disruption of complex networks and was associated with the up upregulation of vascular endothelial growth factor (VEGFA) and down-regulation in vascular endothelial cadherin (VE-Cadherin).

**Conclusion:** These results support, within a biomimetic 3D, in vitro framework, the direct role of CAFs in promoting cancer invasion and that CAFs are also key components in driving vasculogenesis and angiogenesis.

## Cancer associated fibroblasts and tumour growth

The permissive role of the tumour microenvironment in contributing to the process of tumour progression is increasingly recognised^1,2^. Within this complex and dynamic stromal response, cancer associated fibroblasts (CAFs) are of particular interest^3^. CAFs are highly differentiated and activated fibroblasts that comprise a range of subtypes and phenotypes^4^. In the healthy colon tissue, resting fibroblasts line the lamina propia adjacent to the epithelium and precryptal fibroblasts contour the walls of the crypts contributing to tissue integrity^5^. Some subtypes of CAFs are derived from these local fibroblast populations that appear to reside in the margins of the tumour. Other subtypes may migrate from distant sites such as the bone marrow (BM) whilst other are speculated to have derived from other cell types that differentiate into CAFs. CAFs are also believed to be (differentiated) cancer cells through the endothelial/epithelial-mesenchymal transition (EMT)^4,5^. Furthermore, mesenchymal stem cells (MSCs) have been thought to be able to differentiate into CAFs and consequently give rise to other stromal cells such as endothelial cells (ECs)^3^. CAFs promote tumour growth^6^ by the overexpression of growth factors, cytokines, chemokines and matrix-remodelling enzymes whilst increasing stiffness of the tumour^7^. This stiffening in itself can drive tumour growth. Recent work has highlighted the role of stiff tumour tissue on cellular communication network factor 1 (CCN1) regulation in endothelial cells, which enhances melanoma cell-endothelium interaction to promote metastasis through the vasculature^8^. The reactive stroma is in an inflammatory state and under constant stress such as oxygen and nutrient deprivation. CAFs induce the tumour macrophage polarization towards the M2 phenotype, also known as tumour activated macrophages (TAM)^4^, major orchestrators of cancer-related inflammation^9^. This process is driven mainly by interleukin-6 (IL-6), which is highly expressed by CAFs^10^. The key signature of CAFs is the overexpression of alpha smooth muscle actin (αSMA), a contractile stress fibre also expressed by myofibroblasts during wound healing^11^.

A number of “CAF markers” are used to differentiate between normal fibroblasts (NFs) and CAFs. They include fibroblast-specific protein-1 (FSP-1/S100A4), fibroblast-activating protein (FAP), platelet-derived growth factor receptor (PDGFR) and prolyl 4-hydroxylase subunit alpha-1 (P4HA1), however, CAFs are a highly heterogenous population with various activation states present, which makes them difficult to be chracterised^4,7^. Initially, CAFs repress tumour growth due to gap junction formation amongst activated fibroblasts, but consequently they pave the way for extracellular matrix (ECM) remodelling and stiffening^12^. The ECM is remodelled physiologically and chemically during cancer progression due to factors expressed and released by the cancer cells and CAFs, including proteases breaking down the ECM through increased covalent cross-linking of collagen fibrils mediated by lysyl oxidase (LOX)^13,3^. This in turn increases interstitial fluid pressure within the tissue, which activates CAFs to upregulate transforming growth factor beta (TGF-β-1)^4^ and matrix metallopeptidases (MMPs) thus promoting and guiding cancer cell tissue invasion^14^. Stiffness plays a major part in cancer progression and mechanotransduction and YAP-dependent remodelling of the matrix is required for the creation and maintenance of CAFs^15,16^. CAFs produce and excrete a number of soluble factors which stimulate neighbouring stromal cells to excrete further tumour growth supporting soluble factors^17^. This cancer-stroma cross-talk recruits immune cells and local vasculature due to CAFs increasingly excreting vascular endothelial growth factor (VEGF)^10^. Overexpression of IL-6 by CRC cells and CAFs drives cytokinetic angiogenesis and further upregulates VEGF secretion through prostaglandin-E2 (PGE-2) mediation^14^.The recruited vascular networks promote cancer escape from the primary tumour and metastases. Colon CAFs specifically secrete growth factors, like hepatocyte growth factor (HGF), which activates mitogen-activated protein kinase (MAPK) and phosphophatidylinositol 3-kinase (PI3K)/AKT pathways responsible for cell survival and invasion of the cancer^18^.

## CAFs in 3D cancer models

The use of CAFs in *in vitro* 2D and 3D cancer models has been very limited in CRC and using patient-derived samples. CAFs cultured in collagen have increased contractility compared to NFs^13^ and differentiate into cancer cells within co-culture^19^. Previous approaches have used spheroid formation, basic 2D invasion assays and microfluidic devices^20^ in order to replicate the tumour stroma. These approaches are limited in their 3D representation of the tumour stroma by lacking vital components, such as vasculature and a clearly defined tumour-stroma margin through the compartmentalisation of cancer mass and stroma. By replicating the tumour-stroma margin it is possible to study the interplay of different cell populations during cancer progression.

Our approach to engineering a 3D *in vitro* colorectal cancer model incorporates patient-derived CAFs in the stromal compartment and allows us to study the patient-specific effect on vasculature formation during cancer growth and progression. This novel approach of modelling cancer-CAF interplay allows us to directly demonstrate the cellular cross-talk between the cancer and stromal cells within a stable and stiff ECM.

We hypothesised that invasion of cancer cells intro the stromal compartment is enhanced in the presence of CAFs as compared to normal human dermal fibroblasts (HDFs), our control used for this project. We also studied how the presence of CAFs, and the release of growth factors and cytokines, altered the formation of vascular networks and remodelled pre-existing vascular networks.

## Methods

### CAF isolation and propagation

Primary human colorectal cancer associated fibroblasts were isolated from tumour tissues acquired from surgeries at the Royal Free Hospital. Patients provided informed consent for tissue donation for research, ethics code: 11/WA/0077. Fresh samples were provided by the pathology team, ensuring diagnostic margins were not compromised. Tissue was disaggregated using a tumour dissociation kit (Miltenyi Biotec, Bergisch Gladbach, Germany) and grown in Fibroblast Growth Medium 2 (Promocell, Heidelberg, Germany). For the first 72 hours (h) cells were left undisturbed, following that, media changes were done every 48 h in order to isolate the fibroblast cell population. The tissue samples were called T7, T10 and T11 for the first round of successful samples and T6, T9 and T13 for the second lot of successful samples cultured. Patient-derived CAF samples were then tested for positive vimentin expression and negative CK20 expression, to exclude colorectal epithelial cell contamination, and CD31, to eliminate endothelial cell contamination. CAFs αSMA and metabolic activity was assessed (see supplementary data). A range of other general fibroblast and more specific CAF gene markers were also investigated (see *Figure 1B*).

**Figure 1:**
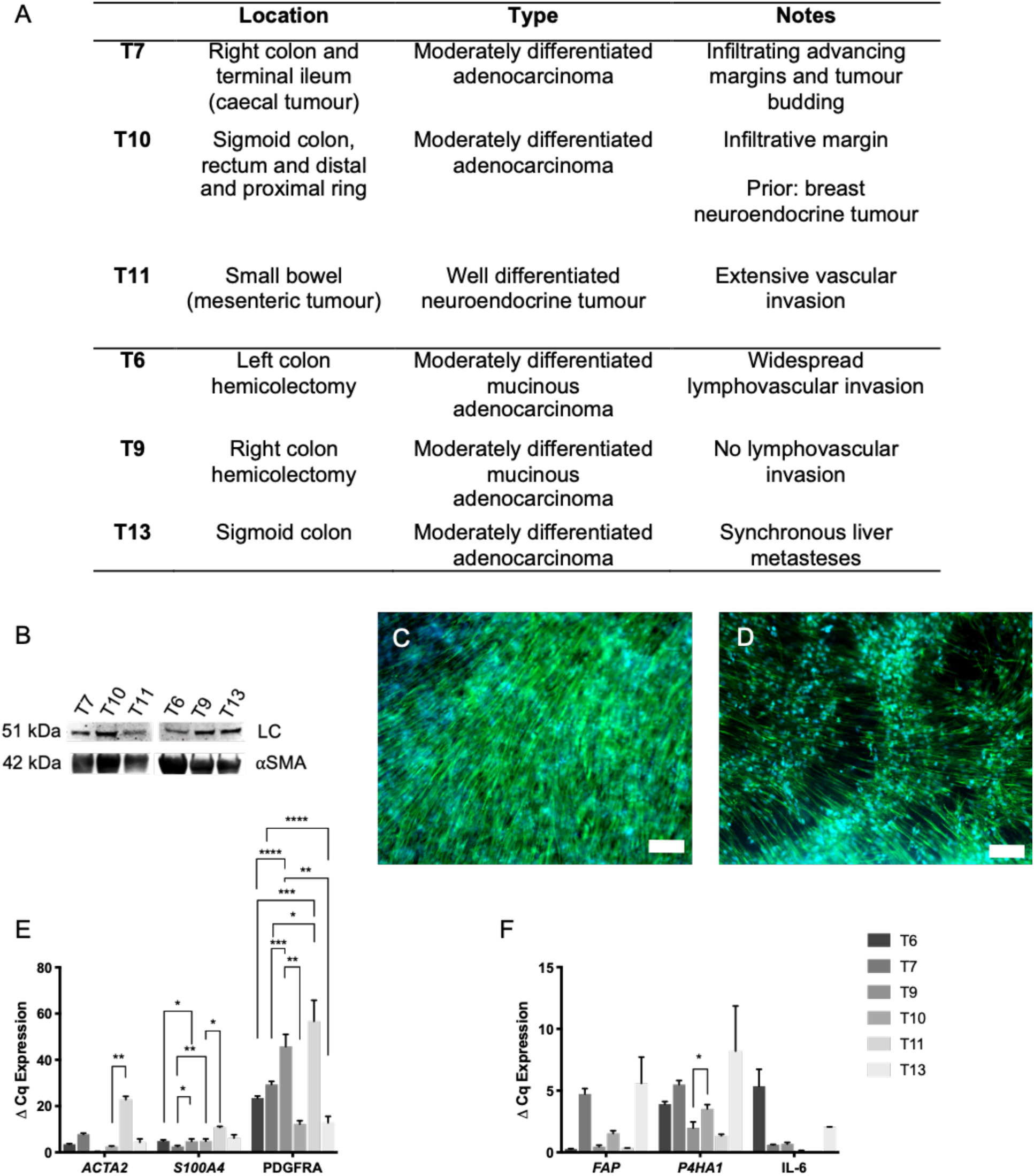
Patient specific CAF tissue sample characterisation. (A) Origin of samples including the location of the original cancer mass, tumour type and any additional notes. (B) Western blot of α-SMA protein expression within 2D monolayers of cells with LC=loading control beta-tubulin. (C) and (D) Visualisation of vimentin expression of HDF and CAFs respectively when cultured in 3D with scale bar=100 μm for both images and green=vimentin and blue=DAPI. (E) 2D cell samples were analysed for fibroblast markers α-SMA (alpha smooth muscle actin), S100A4 (fibroblast specific protein-1), PDGFRA (platelet derived growth factor receptor a) and (F) FAP (fibroblast-activating protein), P4HA1 (prolyl-4 hydroxylase) and IL-6 (interleukin 6). Value shown is normalised to HPRT1 mRNA levels (mean ± SEM) with n=3 and 3 technical repeats. One-way ANOVA with Dunnet’s Post Hoc for ACTA2, S100A4, FAP, PDGFRA and IL-6 and Kruskal-Wallis with Dunn’s Multiple comparisons test p-values for P4HA1, with values 0.05=*, 0.005=**, 0.0005=*** and 0.00005=**** with DOF for first five genes all=20 and f-value for ACTA2=92.1, S100A4=136.7, FAP=31.83, PDGFRA=16.2 and IL-6=13.76.

### Cell culture

Human colorectal adenocarcinoma cell lines HT29 and HCT116 (both European Collection of Cell Cultures through Sigma-Aldrich, Dorset, UK) were grown in Dulbecco’s Modified Eagle Medium (DMEM) at 1 000 mg/L glucose (Sigma-Aldrich, Dorset, UK). Human adult-donor dermal fibroblasts (HDF) (Promocell, Heidelberg, Germany) were grown in 4 500 mg/L DMEM (Sigma-Aldrich, Dorset, UK). Human umbilical vein endothelial cells (HUVEC) were grown in Endothelial Cell Growth Medium (both Promocell, Heidelberg, Germany). After isolation, CAF cells were cultured using Fibroblast Growth Medium 2 (Promocell, Heidelberg, Germany). All media were supplemented with 10% Foetal Calf Serum (FCS) (First Link, Birmingham, UK) as well as 100 units/mL penicillin and 100 μg/mL streptomycin (Gibco™ through Thermo Fisher Scientific, Loughborough, UK). All cell types were cultured at 5% carbon dioxide (CO_2_) atmospheric pressure and at 37°C temperature and routinely passaged in 2D monolayers. HDF and HUVECs were used at passage ≤5.

### Complex 3D models of cancer (tumouroids)

All tumouroids were fabricated using monomeric Type I rat-tail collagen (First Link, Birmingham, UK) and the <u>RAFT™ protocol</u> pages 8-9 (Lonza, Slough, UK) as previously described^21^. 10X MEM (Sigma-Aldrich, Dorset, UK Sigma-Aldrich, Dorset, UK) was mixed with collagen and neutralising agent (N.A.) (17% 10 M NaOH (Sigma-Aldrich, Dorset, UK) in 1 M HEPES buffer Gibco™through Thermo Fisher Scientific, Loughborough, UK)) and mixed with cell suspension resulting in 80% collagen, 10% 10X MEM, 6% N.A. and 4% cells. For the artificial cancer masses (ACMs) 5×10^4^ cells/ACM of either less-invasive HT29 or highly-invasive HCT116 cells were used and 240 μL of the collagen mix was added to a 96-well plate (Corning® Costar® through Sigma-Aldrich, Dorset, UK). The gel mix was polymerised at 37°C for 15 minutes (min), followed by plastic-compression using the 96-well RAFT™ absorbers (Lonza, Slough, UK). In order to produce ‘tumouroids’^22^, the ACMs were nested into a stroma. For the stroma, collagen solution as described above was prepared, and ACMs were directly embedded into a 24-well plate (Corning® Costar® through Sigma-Aldrich, Dorset, UK) containing 1.3 mL of the non-cross-linked collagen mix. Extracellular matrix components were added to this stroma. In this case mouse laminin^23^ 50 μg/mL (Corning® through Sigma-Aldrich, Dorset, UK) for an acellular stroma additionally to 2.5×10^4^ HDFs/CAF samples and 10^5^ HUVECs for a healthy or cancerous stroma respectively (please refer to *Schematic 1* for more detail). The tumouroids were polymerised at 37°C for 15 min and plastic-compressed using the 24-well RAFT™ absorbers (Lonza, Slough, UK). Tumouroids were cultured for up to 21 days at 5% CO_2_ atmospheric pressure and 37°C with 50% media changes every 48 h.

### CAF treatment

To study the effect of CAFs on established endothelial networks, a 1.0 mL suspension containing 2.5×10^4^ CAF cells was added to the media mix at day 21 of established tumouroids. CAFs and tumouroids were subsequently left to propagate in co-culture for 7 days with continuing 48 h 50% media changes. This is additionally demonstrated in the *Schematic 2* below. Investigative measurements were taken at day 21+1, day 21+3 and day 21+7 post CAF addition. *ACTA2* levels were assessed after CAF additiona as an internal control (supplementary section).

### Immunofluorescence

Tumouroids were formalin fixed using 10% neutrally buffered formalin (Genta Medical, York, UK) for 30 min and then washed and stored in phosphate buffered saline (PBS) (Gibco™ through Thermo Fisher Scientific, Loughborough, UK). The tumouroids were permeabilised and blocked for 1 h at room temperature using a solution of 0.2% Triton X 100 and 1% bovine serum albumin (BSA) (both Sigma-Aldrich, Dorset, UK) in PBS. Primary antibody incubation was performed overnight at 4°C followed by three 5 min wash steps with PBS. Secondary antibody incubation was carried out the next day with a 2.5 h incubation at room temperature followed by three 15 min wash steps with PBS. Antibodies were diluted in the same Triton X 100 and BSA solution and suppliers and source were: primary 1:200 anti-CK20 rabbit D9Z1Z (New England Biolabs, Herts, UK), anti-CD31 mouse JC70/A (Abcam, Cambridge, UK) anti-Vimentin mouse V9 (Santa Cruz, Texas, US) and secondary 1:1000 anti-mouse Alexa Fluor® 488 IgG H&L ab150113 and anti-rabbit DyLight® 594 ab96885 (both Abcam, Cambridge, UK). All tumouroids were counterstained with DAPI, using NucBlue™ (Invitrogen™ through Thermo Fisher, Loughborough, UK).

### Measurement of invasion, endothelial networks and analysis

All tumouroids were imaged using the Zeiss AxioObserver with ApoTome.2 and Zeiss ZEN software (Zeiss, Oberkochen, Germany). In order to measure the invasion from the original ACM into the stromal compartment and the number of endothelial structures, 4 images were taken at a 10x magnification evenly spaced out in alignment with a clock face at 12, 3, 6 and 9 o’clock on the same focal plane. This method has previously been described^21^. All samples were assessed for distance and surface area of invasion and number of endothelial structures in the stromal compartment. The images obtained were then analysed in Fiji ImageJ software^24^.

### RNA extraction, cDNa Synthesis and real-time PCR

RNA was extracted using the phase separation TRI Reagent® and chloroform method^25^ (both Sigma-Aldrich, Dorset, UK). Total RNA obtained was quantified and assessed for integrity using the NanoDrop™. Transcription into cDNA was conducted using the High-Capacity cDNA Reverse Transcription Kit (Applied Biosystems™ through Fisher Scientific, Loughborough, UK) on the T100™ Thermal Cycler (Bio-Rad, Watford, UK). Primers were designed according to the MIQE with an annealing temperature (Ta) of 60°C, sequences and efficiencies are listed in *Table 1* below, and were purchased through Eurofins Genomics (Ebersberg, Germany). Gene target amplification was conducted using iTaq™ Universal SYBR® Green Supermix on the CFX96™ Touch System (both Bio-Rad, Watford, UK) in 10μL reactions with 20 ng sample cDNA and primer concentration of 0.2 μM. Relative gene expression was calculated using the ΔCt and 2^-ΔΔCt^ method^26^ normalising to reference gene *hypoxanthine-guanine phosphoribosyltransferase (HPRT1)* with primers for this gene taken from literature^27^. Primer design parameters can be found in the supplementary section. **ELISA.** Media aliquots from cultured tumouroids were taken at every 48 h media change, kept in-80 °C and analysed for vascular endothelia cadherin (VE-Cadherin) active protein expression using the R&D Systems (Abingdon, UK) Human VE-Cadherin Quantikine ELISA Kit according to the manufacturer’s instructions. Results were read on the Tecan Microplate Reader (Männedorf, Switzerland).

**Table 1:**
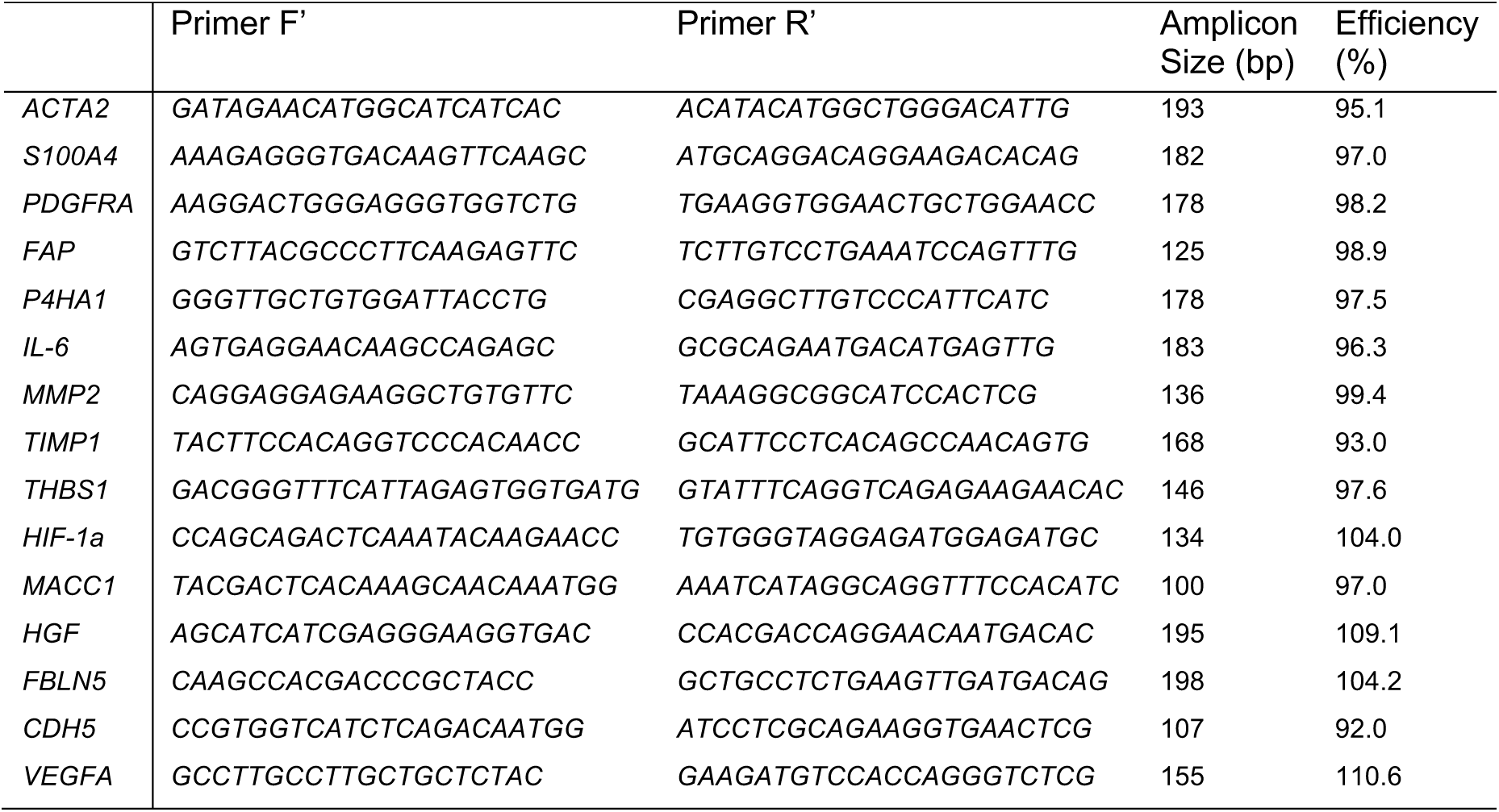
Primer pair sequences, amplicon sizes and efficiencies.

### Protein extraction and western blotting

CAF cell monolayers were lysed for protein with RIPA buffer containing protease inhibitor cocktail at 1:100 dilution (both Sigma-Aldrich, Dorset, UK). Lysed cell suspensions were quantified for protein concentration using the Pierce™ BCA Protein Assay Kit (Thermo Fisher Scientific, Loughborough, UK). Working solutions were made up to 0.5 μg/μl with RIPA and 2x Concentrate Laemmli Sample Buffer (Sigma-Aldrich, Dorset, UK) and 10 μg per sample were loaded onto 10% Mini-PROTEAN^®^ TGX™ Precast 10-well protein gels and run at 200 Volts (V) for approximately 45 min using the Mini-PROTEAN Tetra Cell and PowerPac™ 300 using tris-glycine SDS running buffer (all Bio-Rad, Watford, UK). Protein ladder SeeBlue™ Plus 2 Pre-stained Protein Standard (Invitrogen™ through Thermo Fisher Scientific, Loughborough, UK) was used. Proteins on the gels were then dry transferred onto nitrocellulose membranes using Trans-Blot^®^ Mini Nitrocellulose Transfer Packs and the Trans-Blot^®^ Turbo™ Transfer System (Bio-Rad, Watford, UK). Membranes were blocked for 1 h with 5% milk (Sigma-Aldrich, Dorset, UK) (in tris-buffered saline and 1% Tween 20 (TBST), both Bio-Rad, Watford, UK)), incubated with 1° antibodies for α-SMA 1A4 and loading control β-tubulin N-20 in 5% milk overnight at 4°C at dilutions 1:1000 and 1:200 respectively followed by five quick rinses and three 5 min washed with TBST. 2° antibodies IgG-HRP anti-goat sc-2953 and anti-mouse sc-2314 at 1:1000 dilutions were incubated for 1 h in 3% milk (all antibodies through Santa Cruz Biotechnology, Dallas, US), followed by three 15 min washes with TBST, and developed using Pierce™ ECL Western Blotting Substrate (Thermo Fisher Scientific, Loughborough, UK). Blots were imaged using the ChemiDoc™ XRS imaging system and Image Lab™ software (Bio-Rad, Watford, UK).

### Statistical analyses

All statistical analysis was conducted using GraphPad Prism 7 software. Data was tested for normality with the Shapiro-Wilk test (n≥3) or the D’Agostino test (n≥8) and the appropriate test for statistical significance was applied depending on data parameters (t-test, Mann-Whitney, One-way ANOVA with Dunnet’s Post Hoc or Kruskal-Wallis with Dunn’s multiple comparisons test). The tests used for each graph are outlined within the figure legends individually. Significance was at p-values <0.05. All data points are represented as mean with standard error mean (SEM) in graphs and values stated in text as mean with standard deviation (STDEV). In general, n=3 with 3-4 technical replicates, details described within the figure legends for each individual data set. F-values, t-values and degrees of freedom (DOF) are stated within the figure legends for each set of statistical tests. Two-tailed tests for significance were used when appropriate.

**Schematic 1:**
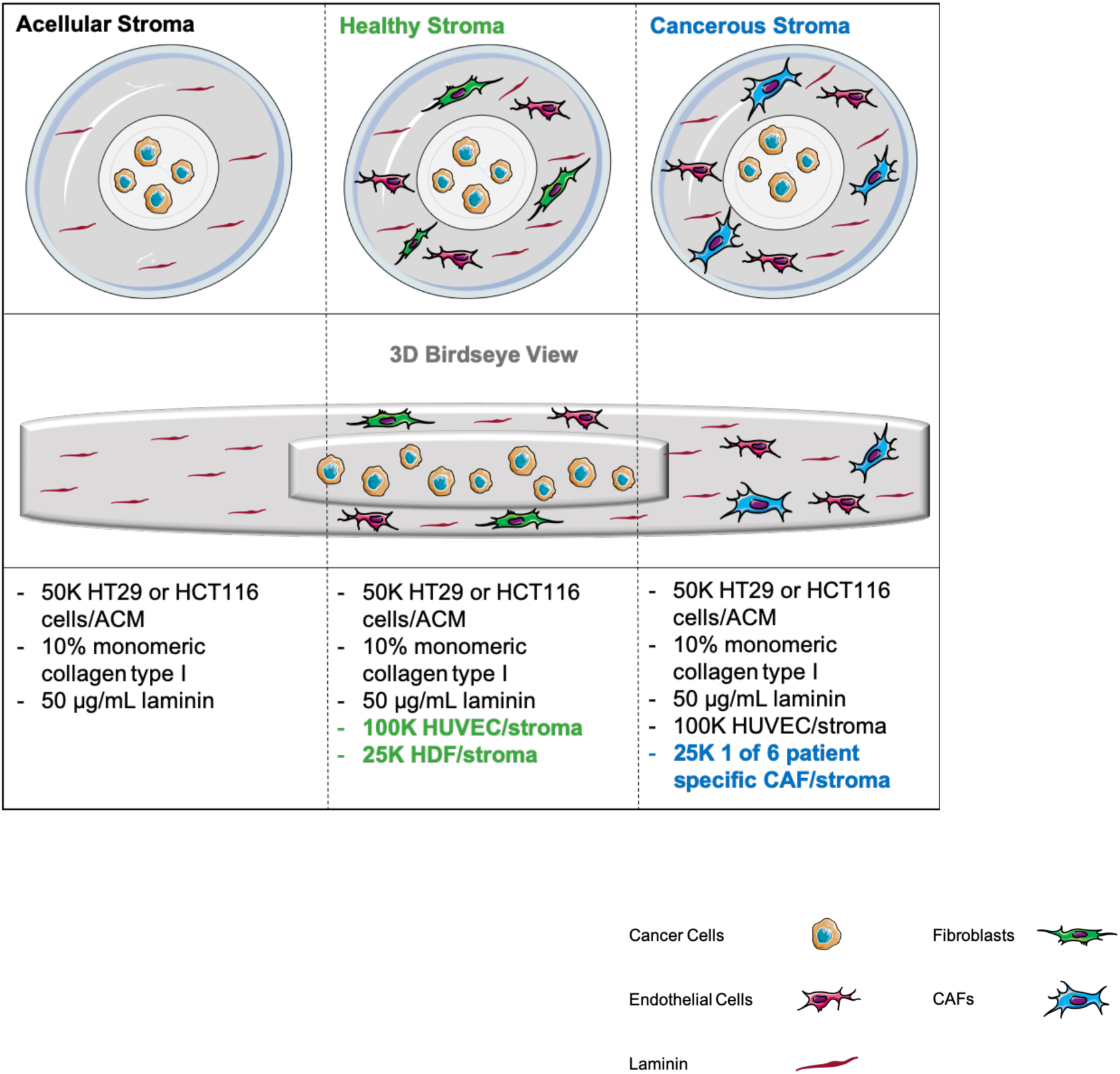
The three main tumouroid set ups used with respective cellular populations in the ACM and stroma. For all set ups, an ACM was nested into a stromal compartment. Both consisted of 10% monomeric collagen type 1 that had undergone plastic compression with the RAFT™ protocol. Schematic was created using Servier Medical Art according to a Creative Commons Attribution 3.0 Unported License guidelines 3.0 (https://creativecommons.org/licenses/by/3.0/).

**Schematic 2:**
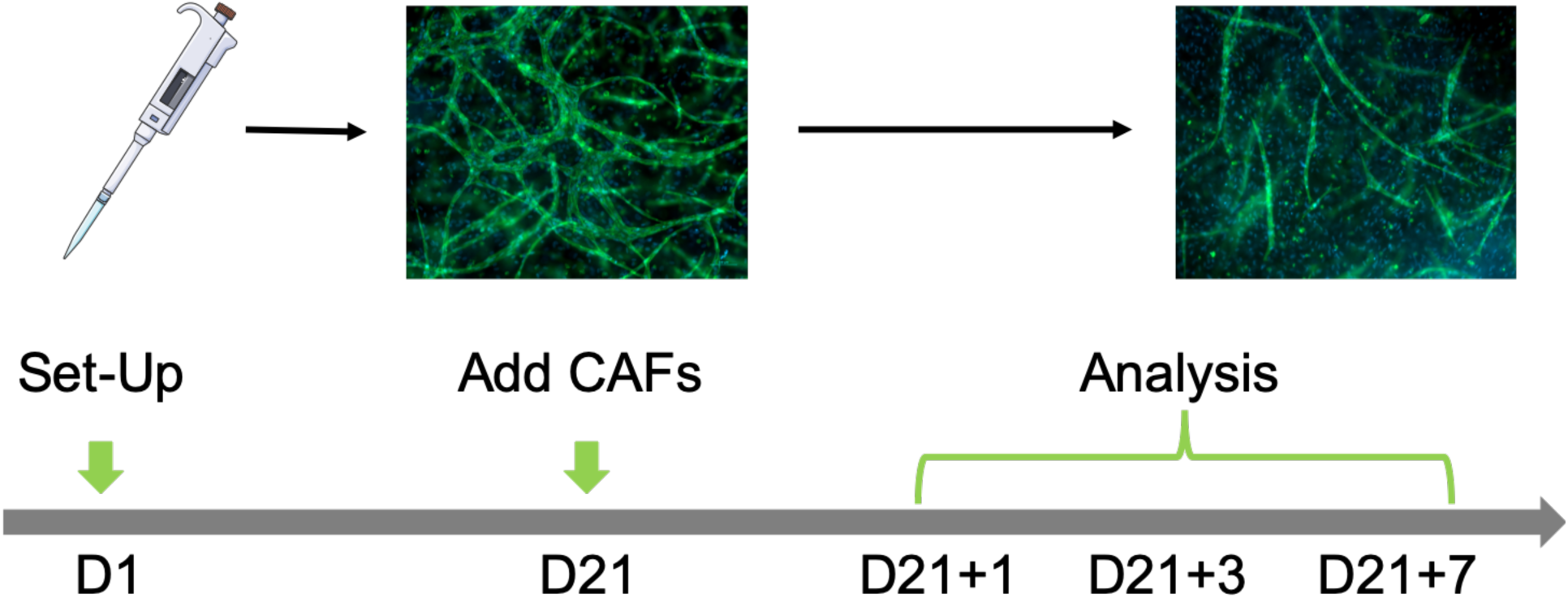
The “CAF Treatment” set-up. Tumouroids with a normal HDF containing stroma were left to mature for 21 days. During this time endothelial networks developed. After this time, one of three patient-specific CAF samples were applied to the mature tumouroids. This resulted

## Results

### Extraction, propagation and characterisation of patient-derived CAF samples

Six patient-derived CAF samples (n=6) were established from tumour samples, expanded on 2D tissue culture plastic (passage ≤3) and included in the tumouroid model. The samples were of variable location and origin (*Figure 1A*), but all samples were from the lower bowel, colon or rectum with 5 being of adenocarcinoma and 1 being of neuroendocrine type. Samples were of varying grade in terms of tumour margin infiltration and vascular invasion. All samples were successfully cultured in 2D monolayers and tested for a number of fibroblast markers at the gene level (*Figure 1E&F*). The data showed that all six samples were positive for *ACTA2, S100A4, PDGFRA, FAP, P4HA1* and *IL-6*. This confirms that the cells are activated fibroblasts, especially based on the high expression of *S100A4, PDGFRA* and *IL-6* in all samples^28,29&30^. Gene expression levels were compared between the samples and their differences can be taken from *Figure 1E&F*. Secondly, the western blots showed that the α-SMA protein was expressed in all samples (*Figure 1B*), this is a measure previously used to distinguish samples as CAFs^31^. Thirdly, vimentin staining was done in CAF tumouroids grown to confluency and the morphology was compared to normal HDFs within tumouroids (*Figure 1 C&D*). It was observed that the CAF samples overall appeared to have more stress fibres and had a much less organised structure.

### A healthy stroma does not upregulate cancer invasion significantly

There is an evident cross-talk between the cancer cells and surrounding stroma. In the tumouroids, extensive primitive endothelial networks are formed within the stroma whilst the fibroblast population completely overgrows the space (*Figure 2D,E&F*).

**Figure 2:**
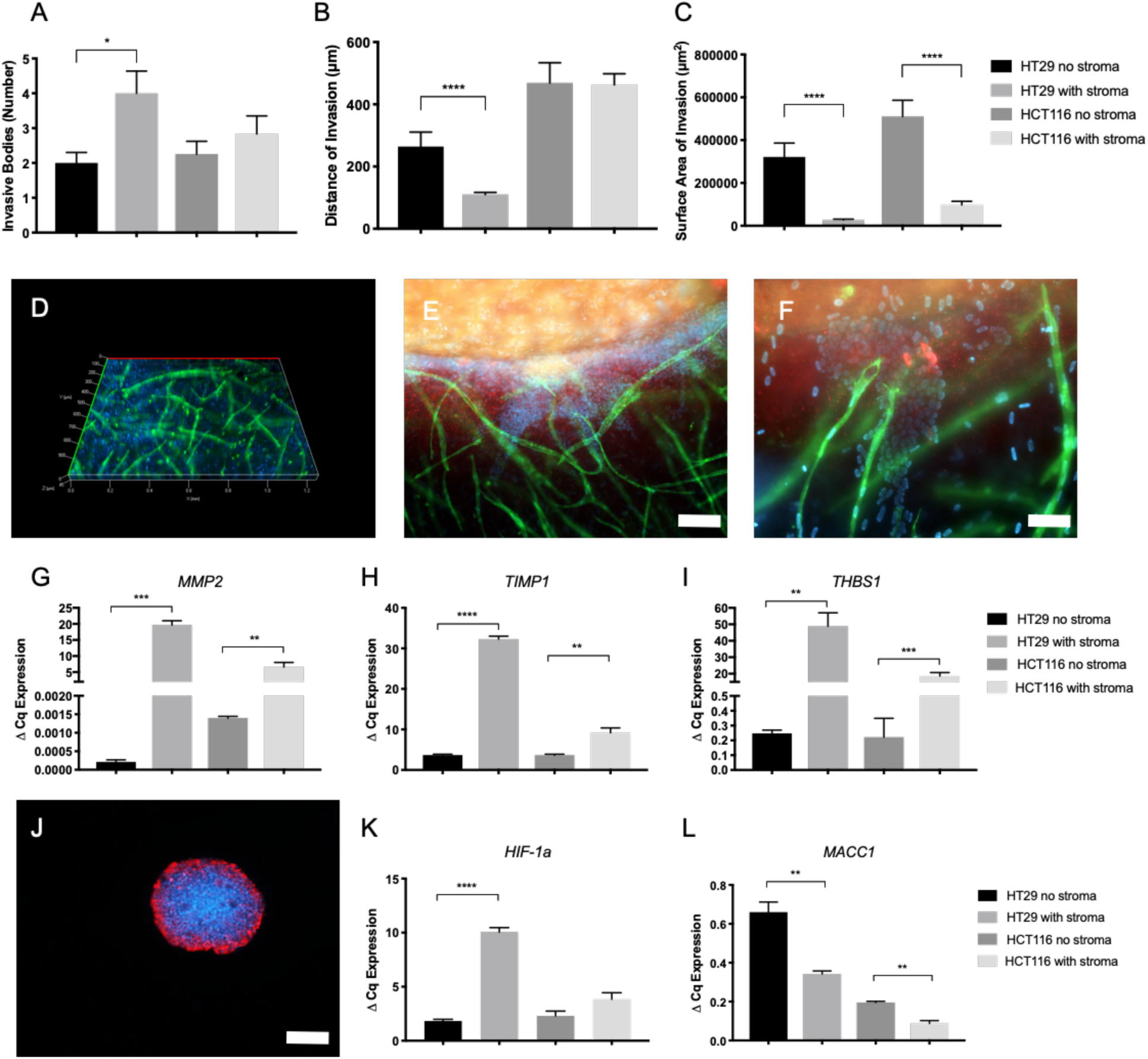
Invasion into an acellular or healthy cellular stroma. (A) Number of invasive bodies, (B) distance of invasion and (C) average surface area of invasion at day 21 of HT29 or HCT116 toumoroids (mean ± SEM). All n=3 with 4 technical repeats and showing Mann-Whitney p-values, with values 0.05=*, 0.005=**, 0.0005=*** and 0.00005=****. (D) Representative z-stack of vascularized/cellular stroma, (E) invasion into cellular stroma within HCT116 tumouroids and (F) interaction between cancer cells and endothelial structures with scale bar=100 μm and 50 μm respectively and with red=CK20, green=CD31 and blue=DAPI. Comparative gene expression between acellular and cellular stroma in tumouroids at day 21 of growth for (G) matrix metallopeptidase 2 (MMP2), (H) metallopeptiase inhibitor 1(TIMP-1), (I) thrombospondin 1 (THBS1), (K) hypoxia inducible factor 1-alpha (HIF-1a) and (L) metastasis associated in colon cancer-1 (MACC1).Value shown is normalised to HPRT1 mRNA levels (mean ± SEM) with n=3 and 3 technical repeats showing Unpaired t-test p-values, with values 0.05=*, 0.005=**, 0.0005=*** and 0.00005=****. (J) Representative image of HT29 invasive body at day 21, not containing a cellular stroma showing a large surface area with scale bar=100 with red=CK20, and blue=DAPI.

The effect of adding cells to the stromal compartment was measured comparing tumouroids with an acellular stroma to ones containing normal fibroblasts and a primitive vascular network within the stroma. Firstly, the number of invasive bodies increased in the presence of a cellular stroma (*Figure 2A*), significantly in the HT29 tumouroids (p=0.0123) with an average of 2.000±1.044 in an acellular stroma compared to 4.000±2.216 average invasive bodies per image analysed. However, the average distance of invasion decreased significantly (*Figure 2B*) in the HT29 tumouroids (p=0.0006) with an average distance of invasion within the acellular stroma of 264.3 μm±138.9 μm and 110.2 μm± 47.7 μm. The surface area of invasion also decreased significantly (*Figure 2C*) in the presence of a cellular stroma (p<0.0001) for both the HT29 and HCT116 tumouroids. For the HT29 acellular stroma the average surface area of invasion was 321,801 μm^2^± 182,232 μm^2^ and 26,963 μm^2^±26,871 μm^2^ within a cellular stroma and within the HCT116 tumouroids it was 510,240 μm^2^±313,463 μm^2^ for acellular and 97,946 μm^2^± 95,469 μm^2^ for a cellular stroma present. This means that although the number of invasive bodies increased when incorporating a cellular stroma, the distance and surface area invaded decreased (*Figure 2J*).

When analysing the genes responsible for the different invasion patterns, a number of genes were significantly upregulated when going from an acellular to a cellular stroma including *MMP2* for HT29 and HCT116 tumouroids (p=0.001 and p=0.0098 respectively), *TIMP1* (p=<0.0001 and 0.0099 respectively) and *THBS1* (p=0.0039 and p=0.0007 respectively) (*Figure 2G, H&I*). Additionally, *HIF-1a* was upregulated (*Figure 2K*) in the presence of a cellular stroma compared to an acellular stroma in HT29 tumouroids (p=<0.0001), indicating that there was more hypoxia occurring within the tumouroids. Interestingly, *MACC1* was downregulated in the presence of a cellular stroma within the HT29 and HCT116 tumouroids (p=0.0041 and 0.0024 respectively) (*Figure 2L*).

### A cancerous stroma significantly upregulates cancer invasion

CAFs were incorporated into the cancer stroma within the 3D tumouroid model containing either a HT29 or HCT116 ACM in order to investigate the effect of a cancerous stroma on cancer growth. The CAF-derived stroma caused an increase in the distance and surface area of invasion compared to HDF-derived stroma (*Figure 3A,B,E&F*). For the less-invasive HT29^32^ tumouroids, samples T6, T10, T11 and T13 caused a significant upregulation in the distance of invasion (p=<0.0001, 0.0014, <0.0001 and <0.0001 respectively) with an average distance of invasion (μm) as follows: T6=320.4±162.3, T10=239.9±132.3, T11=352.4±209.9 and T13=361.9±224 as compared to HDF containing tumouroids =110.2±47.7. In the highly-invasive HCT116 tumouroids, the CAFs statistically increased the average distance of invasion (μm) in the presence of sample T13 (p=0.0489) =693.6±253.2 compared to HDF containing =463±207. The average surface area (μm^2^) invaded for the HT29 tumouroids was significantly greater in the presence of samples T6, T11 and T13 (p=<0.0001 for all three) with T6=205,140±164,304, T11=290,460±311,530 and T13=203,120±193493 compared to HDF containing =26,963±26,871. For the HCT116 tumouroids, the average surface area invaded (μm^2^) by the cancer was significantly upregulated in the presence of samples T6, T11 and T13 also (p=<0.0001 for all three) with T6=415,796±181,028, T11=456,344±232,945 and T13=537,545±257466 as opposed to HDF containing being =97,946±95,469. This is shown in the images taken by day 21 of tumouroids (*Figure 3C&D*), which demonstrate the increase in size of the invasive bodies in the presence of CAFs. A panel of 30 genes involved in invasiveness and angiogenesis were investigated to compare the healthy and cancerous stroma. Genes that were significantly upregulated in CAF-tumouroids were *HGF, ACTA2* and *TIMP1 (Figure 3G,H&I respectively)*. In the HT29 tumouroids, *HGF* was upregulated significantly in the presence of T9 (p=0.0105) and within the HCT116 tumouroids, *HGF* was significantly upregulated in the presence of samples T7 (p=0.0001), T10 (p=0.0071) and T11 (p=0.0255). There was a tendency for increased *ACTA2* in CAF-containing HT29 tumouroids, but it was not statistically significant whilst for the HCT116 tumouroids, *ACTA2* was upregulated significantly in the presence of samples T7 (p=0.0015) and T10 (p=0.0457). Finally, *TIMP1* was highly overexpressed in the presence of CAF samples. In the HT29 tumouroids, the presence of samples T6 (p=0.0010), T9(p=0.0001), T10 (p=0.0026) and T13 (p=0.0001) significantly increased *TIMP1* expression and in the HCT116 tumouroids, *TIMP1* expression was significantly increased in the presence of samples T6 (p=0.0264), T9 (p=0.0329), T10 (p=0.0001) and T13 (p=0.0087).

**Figure 3:**
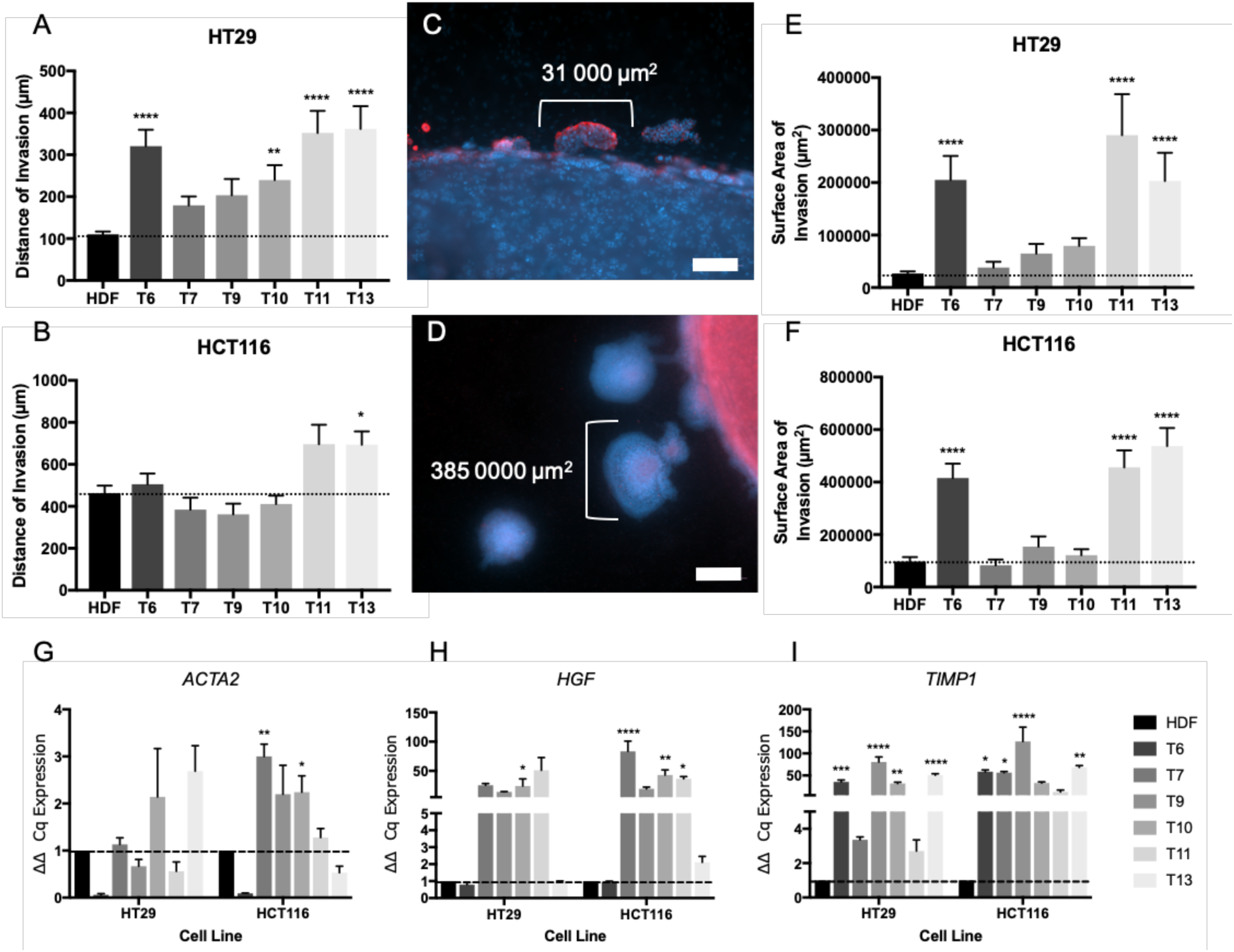
Invasion into stromal compartment and gene upregulation. (A) HT29 distance of invasion and (B) HCT116 distance of invasion into the stroma within tumouroid models at D21 and surface are of invasion within (E) HT29 and (F) HCT116 tumourids at D21. Tumouroids contained either HDF cells or one of six patient-specific CAF tissue samples within the stromal compartment of the constructs. All mean ± SEM with n=3 with 4 technical repeats and showing Kruskal-Wallis with Dunn’s multiple comparison’s test p-values, with values 0.05=*, 0.005=**, 0.0005=*** and 0.00005=****. (C) Representation of average invasive bodies at day 21 in a ‘normal’ HDF containing tumouroid in comparison to a (D) CAF containing tumouroid. Scale bar=100 μm for top and 500 μm for bottom image, with red=CK20 and blue=DAPI. (G) ACTA2 (α-smooth muscle actin), (H) HGF (hepatocyte growth factor and (I) TIMP1 (metallopepdidase inhibitor 1) gene expression in tumouroids at day 21 of growth, comparing HDF containing stroma and CAF containing stroma. Value shown is normalised to HPRT1 mRNA levels (mean ± SEM) with n=3 and 3 technical repeats. Ordinary one-way ANOVA Dunnett’s multiple comparisons test with p-values 0.05=*, 0.005=**, 0.0005=*** and 0.00005=**** with DOF=20 for all and f-value for ACTA2 for HT29=4.213 and HCT116=12.31, f-value for HGF=3.836 for HT29 group and 16.3 for HCT116 group, and finally for TIMP1 f-value for HT29=37.21 and HCT116=11.25.

### The presence of CAFs within the tumouroid stroma inhibits vascular network formation

CAF containing tumouroids demonstrated an inhibition of vasculogenesis; the *de novo* formation of endothelial networks^33^. This was seen as a decrease in the number of elongated endothelial structures formed within the CAF stroma by day 21 of tumouroid culture. In the HT29 tumouroids (*Figure 4A*), the number of endothelial structures was reduced significantly in the presence of samples T6, T7, T11 (all p=<0.0001) and T13 (p=0.0008) with the average number of endothelial structures being T6=0.4167±0.6686, T7&T11=0±0, and T13=2.833±2.25 compared to endothelial structures in HDF containing tumouroids averaging 41.92±5.468. In the HCT116 tumouroids (*Figure 4B*), the presence of CAFs significantly decreased the average number of endothelial structures for samples T6, T9 (both p=<0.0001), T10 (p=0.0479), T11 (p=0.0215) and T13 (p=<0.0001) with average number of endothelial structures T6=0.25±0.4523, T9=5.67±9.664, T10=15.79±27.21, T11=14.86±22.98 and T13=0.179±1.488 compared to HDF-tumouroids=50.67±4.697. Whilst in the HDF containing stroma, endothelial structures formed throughout the entire stromal compartment of the tumouroids (*Figure 4C&D*), in the CAF containing stroma the formation of complex endothelial structures was only observed around invasive bodies from the cancer mass (*Figure 4E&F*). The protein levels of VE-Cadherin, a protein involved in endothelial cell end-to-end fusion^34^, showed a temporal decrease in VE-cadherin levels over the 21 day culture period. The amount of produced VE-Cadherin (ng/mL) significantly decreased in the HT29 tumouroids with sample T13 (p=0.0148) from 22.07±2.144 at day 2 to 12±1 by day 21. Within the HCT116 tumouroids, the VE-Cadherin production significantly decreased in the presence of T6 (p=0.0413) going from 25.04±2.649 at day 2 to 13.86±1.415 by day 21. The gene expression level of *FBLN5*, a gene which inhibits endothelial cell proliferation^35^, angiogenesis^36^ and especially sprouting^37^, were measured in CAF- and HDF-tumouroids at day 21 (*Figure 4I*). In the HT29 tumouroids samples T9, T10 (both p=0.0001) and T13 (p=0.0007) caused a significant upregulation in the relative gene expression while in the HCT116 tumouroids, samples T7 (p=0.0063) and T9 (p=0.0014) significantly upregulated *FBLN5* compared to the HDF-tumouroids.

**Figure 4:**
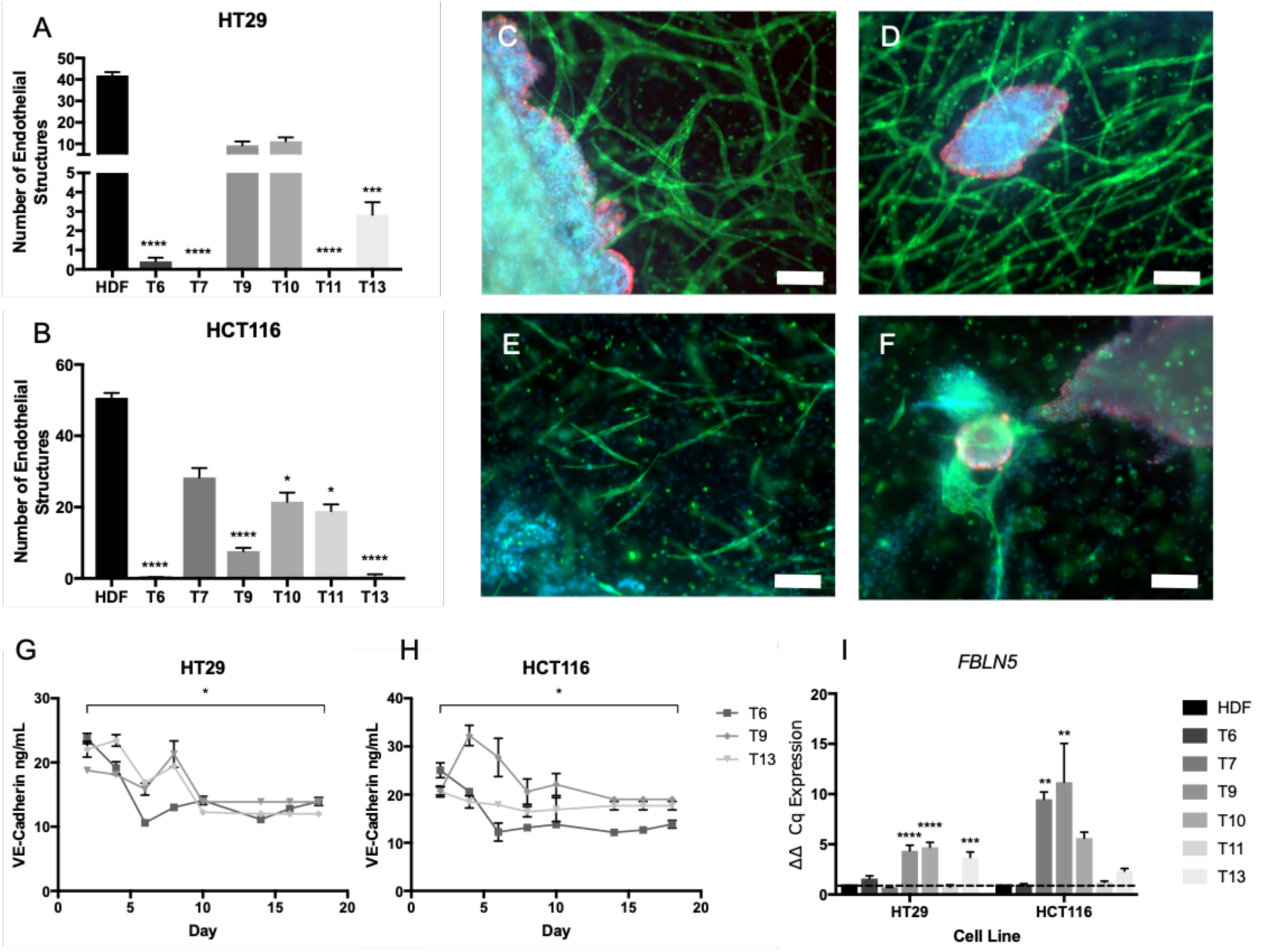
Endothelial structures formed within the cancerous CAF stroma. Number of endothelial structures formed within (A) HT29 and (B) HCT116 tumouroid stromal compartments at day 21 (mean ± SEM) containing either HDF or one of six patient-specific CAF tissue samples. All n=3 with 4 technical repeats and showing Kruskal-Wallis with Dunn’s multiple comparison’s test p-values, with values 0.05=*, 0.005=**, 0.0005=*** and 0.00005=****. (C) Example images of normal endothelial structure formation within a HDF containing stroma at day 21 at the cancer-stromal edge and with a (D) budded invasive body. Scale bar=100 μm for left and 50 μm for right image and red=CK20, green=CD31 and blue=DAPI. (E) Images showing the decreased formation of complex endothelial structures within a CAF containing stroma at day 21 at the cancer-stroma edge, (F) as well as around a budding invasive body, the only occurrence of endothelial structures within these conditions. Scale bar=100 μm for left and 50 μm for right image and red=CK20, green=CD31 and blue=DAPI. Active VE-Cadherin protein released into the media of one of six CAF containing (G) HT29 or (H) HCT116 tumouroids over 21 days (mean ± SEM). Paired t-test comparisons test between day and day 21 with p-values 0.05=*, 0.005=**, 0.0005=*** and 0.00005=**** with DOF=2 for both and t-value for HT29=8.115 and HCT116=4.766. (I) FBLN5 (fibulin-5) expression at day 21 within the HDF or one of six CAF containing HT29 or HCT116 tumouroids (mean±SEM). Value shown is normalised to HPRT1 mRNA levels (mean ± SEM) with n=3 and 3 technical repeats. Ordinary one-way ANOVA Dunnett’s multiple comparisons test with p-values 0.05=*, 0.005=**, 0.0005=*** and 0.00005=****.

### The disruption of pre-formed vascular networks by CAFs

In order to investigate the effect of CAFs on a developed, mature endothelial network (angiogenesis), CAF samples were added on top of tumouroids at day 21 and propagated for 7 days. Endothelial cells start off as single cells within the stroma on day 1 (*Figure 5A*) and form complex, branched networks by day 21 (*Figure 4B*) in the presence of HDFs within a tumouroid. After CAF addition, a disruption of the endothelial networks was observed (*Figure 5C*). The disruption of vascular networks was assessed by quantifying the number and complexity of endothelial structures. The number of endothelial structures on day 21+7 decreased in the presence of all three CAF samples for both the HT29 tumouroids and HCT116 tumouroids (*Figure 5D&E*). For the HT29 tumouroids, samples T6, T9 and T13 significantly decreased the number of endothelial structures after 7 days (p=0.0155, 0.0003 and <0.0001) with the average number of endothelial structures T6=17.42±4.033, T9=15.17±2.623 and T13=9.333±2.741 compared to HDF containing tumouroids with 29±3.384. Within the HCT116 tumouroids, samples T6, T9 and T13 also caused a significant decrease in the average number of endothelial structures after 7 days (p=<0.0001, <0.0001 and 0.0029) with values T6=10.67±3.42, T9=9.083±3.753 and T13=13.08±4.66 compared to HDF containing tumouroids with 27.33±4.942. The vascular disruption was further confirmed by a significant decrease in *CDH5* gene levels, coding for VE-Cadherin. Relative CDH5 levels decreased significantly (p<0.0001 for all) in both HT29 and HCT116 tumouroids at day 21+1, day 21+3 and day 21+7 (*Figure 5F&G*). Furthermore, *FBLN5* gene expression increased significantly after CAF addition (*Figure 4H&I*). In HT29 tumouroids containing samples T6 and T9 on day 21+7 (p=0.0001 and 0.0395). Within the HCT116 tumouroids, *FLBN5* was upregulated significantly for T13 at day 21+1 (p=0.0001), T9 and T13 for day 21+3 (p=0.0261 and 0.0024) and T6 and T13 for day 21+7 (p=0.0300 and 0.0001). Finally, *VEGFA* gene levels were analysed after CAF addition (*Figure 5J&K*) and a general increase was measured. Within the HT29 tumouroids sample T9 caused a significant upregulation at day 21+1 (p=0.02304) and samples T6 and T9 at day 21+7 (p=0.0233 and 0.0072). For the HCT116 tumouroids sample T13 caused a significant upregulation at day 21+1 (p=0.0372), samples T6, T9 and T13 at day 21+3 (p=0.0008, 0.0071 and 0.0019) and sample T13 for day 21+7 (p=0.0282).

**Figure 5:**
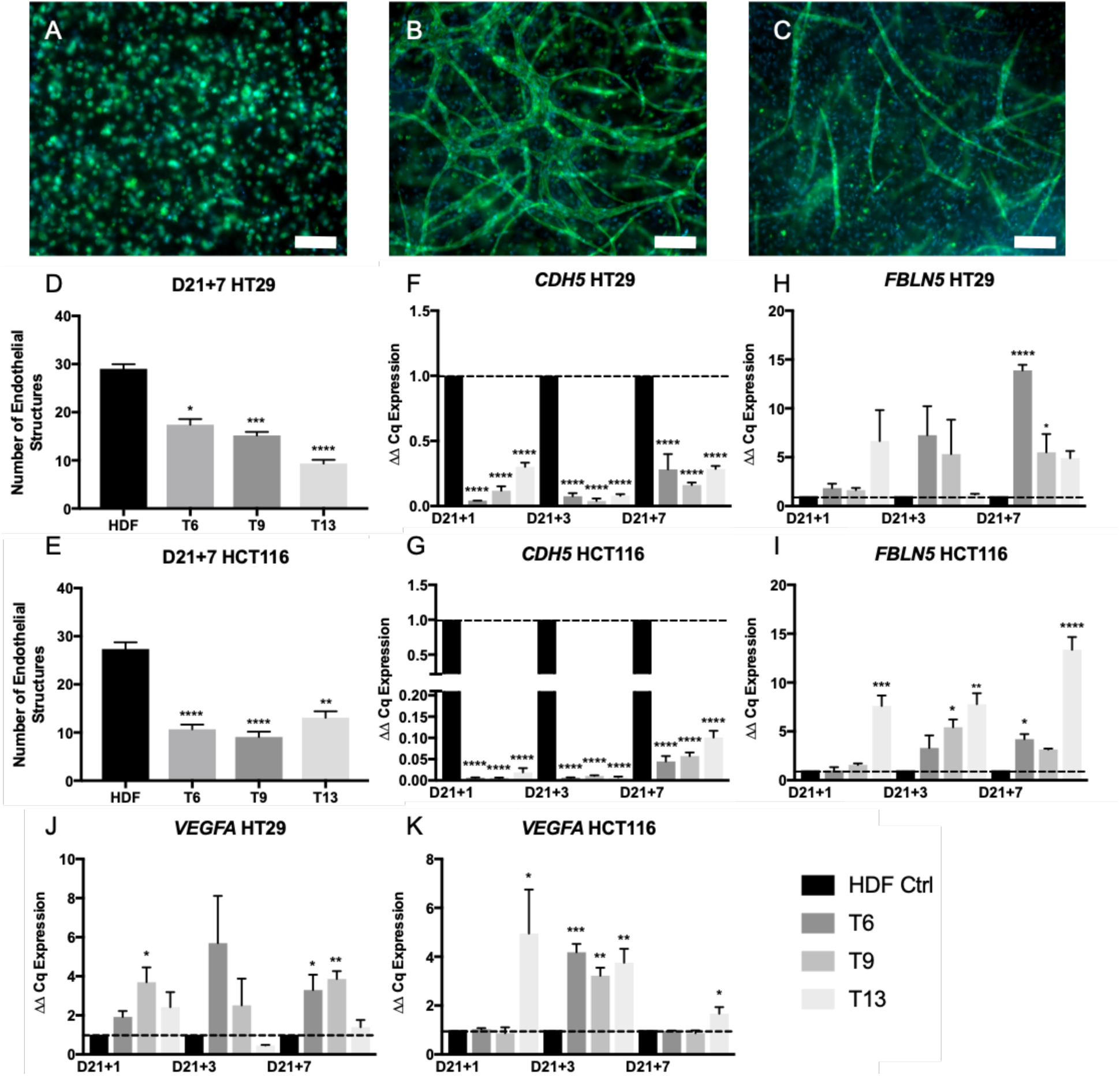
Disruption of mature endothelial network caused by the addition of CAFs. (A) Example of single cell endothelial cells at day 1 of tumouroid growth, (B) example of matured networks at day 21 in an HDF containing stroma and finally (C) example of disrupted networks at day 21+7 (21 days normal/HDF stroma growth and 7 days after CAF addition). Scale bar=100 μm and green=CD31 and blue=DAPI. Number of endothelial structures formed within (D) HT29 and (E) HCT116 tumouroid stromal compartments at day 21+7 (mean ± SEM) containing either HDF or one of six patient-specific CAF tissue samples. All n=3 with 4 technical repeats and showing Kruskal-Wallis with Dunn’s multiple comparison’s test p-values, with values 0.05=*, 0.005=**, 0.0005=*** and 0.00005=****. CDH5 (VE-Cadherin) expression in (F) HT29 and (G) HCT116 tumouroids, FBLN5 (fibulin-5) expression in (H) HT29 and (I) HCT116 tumouroids and VEGFA (vascular endothelial growth factor) expression in (J) HT29 and (K) HCT116 tumouroids after CAF addition at days 1, 3 and 7 (mean±SEM). Value shown is normalised to HPRT1 mRNA levels (mean ± SEM) with n=3 and 3 technical repeats. Ordinary one-way ANOVA Dunnett’s multiple comparisons test with p-values 0.05=*, 0.005=**, 0.0005=*** and 0.00005=****.

## Discussion

Our findings can be summarised as follows. First, a normal healthy stroma does not upregulate cancer growth significantly in a 3D model with a defined cancer mass and defines stromal compartments. Secondly, the presence of a cancerous CAF stroma increased the distance and surface area of invasion of colorectal cancer (CRC) into the stromal compartment whilst, at the same time, inhibiting vasculogenesis. These processes were driven by the up-regulation of hepatocyte growth factor (*HFG*), metallopeptidase inhibitor 1 (*TIMP1*) and fibulin 5 (*FBLN5*). Next, the re-modelling appeared to occur through the process of disruption of complex endothelial networks and was associated with up-regulation of vascular endothelial growth factor (*VEGFA*) and a down-regulation in vascular endothelial cadherin (VE-Cadherin). These results support, within a biomimetic, 3D, *in vitro* framework, the direct role of CAFs in promoting cancer invasion and driving both vasculogenesis and angiogenesis.

The first of our observations describes the effect of adding a healthy stroma to surround the engineered cancer mass. This 3D model, comprising a cancer mass and a stroma, permits the study of how cancer cells invade from a distinct original mass into a new healthy tissue or stroma in the context of a metastasis. Future advancements to this model would include an active immune component. The only aspect that was increased in the presence of a healthy stroma was the increase in invasive bodies, modelling the highly invasive cancers “dispersal” into a high number of small clusters to invade the tissue. This is often due to the loss of structure proteins such as cadherins and cytokeratins^38^. This underlines the necessity of a cancerous storma for a cancer to thrive. The second major finding of this novel work is the differential invasion rate and pattern of less-invasive HT29 and highly-invasive HCT116 cancer cells in 3D tumouroids in the presence of CAFs, causing an increase in the distance (up to 3-fold) and surface area of invasion (up to 10-fold) over a 21-day period. CAFs pro-invasive properties and their ability to enable cancer cells to metastasise has been demonstrated previously^39,40^. Logsdon et al.^41^ described the importance of CAFs in a 3D pancreatic cancer model using Matrigel®-coated invasion chambers and soft-agar colony formation and although this model showed an increased in proliferation and metastasis during *in vivo* validation, the 3D model was not compartmentalised and did not allow for a measurement of invasion *in vitro*. The majority of 3D cancer models lack appropriate tensile force and stiffness associated with tumour tissue, as they commonly use soft hydrogels, which have too high a water content^42^. Our biomimetic 3D *in vitro* cancer model (tumouroid) has a collagen density of up to 40x higher compared to standard hydrogels and therefore mimics the *in vivo* stiff tumour environment more closely^22^; an important aspect especially for CAFs^43^. Our tumouroids let us investigate invasion patterns and study the genes that may be responsible for the increase in invasion. From a personalised healthcare perspective, the prospect of culturing patient specific cells in the presence of a highly or less invasive cancer mass allows the possibility to interrogate the interaction of these two cell populations. Primary cancer cells are especially difficult to culture and therefore utilising patient specific CAFs instead creates an avenue to overcome this limitation and supplement information we can retrieve.

The third of our observations can be interpreted in the following manner; CAFs play a key role in vascular network formation and remodelling. Whilst it is understood that CAFs play a major role in angiogenesis and recruiting vasculature towards the cancer^29^, in this study we also demonstrated that CAFs play a major role in vasculogenesis and the disruption of vascular network formation. This aspect has not been studied with the same rigour. Some studies have introduced CAFs at the same time point as HUVECs and observed end-to-end fusion of the HUVEC cells into endothelial structures^44^. Whilst this could be an observation of vasculogenesis, our model, with tissue specific parameters including biomimetic matrix density, shows no *de novo* formation of vascular networks in 3D in the presence of CAFs. At 25 000 CAFs per 24-well tumouroids, we observed 100% confluency of these cells in 3D by day 7, whilst HUVECs in our “normal” HDF containing cultures would not start forming complex endothelial structures until at least day 14. Our data indicates that CAFs start expressing factors that block complex vascular network formation, whilst retaining ‘simple’ vascular/endothelial structures. One of the factors, that was significantly increased, was *VEGFA.* Although *VEGFA* is the major player in angiogenesis and involved in recruiting mature blood vessels towards the cancer, its role in vasculogenesis is not as well understood. The major cross-talk between cancer cells and endothelial cells in our set up was growth factor driven, which was ascertained through an additional 3D set up. This set up demonstrated the chemoattractant driven movement and recruitment of endothelial structures to the cancer mass through an acellular ring placed between the cancer mass and stromal compartment^21^. Interestingly, a study looking at cardiac mouse development found a correlation between the disruption of vasculogensis and elevated *VEFGA* levels^45^. Within our model we observed the inhibition of vascular network formation in the presence of CAFs and, even within the HDF-tumouroids, we observed that closer to the central cancer mass, the endothelial networks became less complex. This indicates that the additional VEGFA released by the cancer cells could be inhibiting vasculogenesis close to the cancer mass and further underlines the fact that highly metastatic cancer cells and its stromal counterparts may disrupt the initial formation of highly complex endothelial networks. This was further studied by Brown et al. when looking at how the prevention of vasculogenesis but not angiogenesis prevented the recurrence of glioblastoma in mice^46^. Cancer cells are known for their high turnover and over-production of angiogenic growth factors^47^. This is in an attempt to recruit host vasculature from surrounding tissues. The unregulated and upregulated production and release of angogenic growth factors by solid tumours results in the formation of abhorrent and leaky vasculature surrounding tumours^48^. We have measured increased levels of VEGFA in our tumouroid cultures which are resulting in disrupted vasculogenesis and angiogenic remodelling to form non-complex networks. Vasculogenesis in cancer and especially in relation to the presence of CAFs is not a major focus of research as angiogenesis and remodelling of cancer is the biomimetic environment in which cancer arises. However, by gaining insights into how cancer angiogenic signalling can influence vasculogenesis may help further our understanding of tissue necrosis and vascular remodelling in cancer.

In our third observation we further showed that CAFs have the ability to disrupt pre-formed *in vitro* vascular network (“CAF treatment”). By day 7, post CAF addition, the endothelial networks that had previously formed were disrupted and an overall decrease in the number of endothelial structures was observed. Furthermore, we found a decrease of *CDH5* levels within the tumouroids. The role *CDH5* and the corresponding protein VE-Cadherin is becoming more pertinent in the study of angiogenesis as it has been specifically implicated in the local production of junctions within complex endothelial networks^34^. The literature on CAF interaction with VE-Cadherin is limited, although the role of CAFs as major sources of VEGFA production is established and understood to be mediated through HIF-1*α*/GPER signaling^49^. This particular gene (*VEGFA*) was increased after we added CAFs to our cultures and in fact this will have played an important role in the disruption (or angiogenesis) observed, however it would be crucial to study further how stromal cells cause this angiogenesis as the normalisation of these remodelled vascular networks has been a target for many antiangiogenic drugs^50^.

Our results need to be understood in the context of the following methodological limitations. We report the interactions at the interface of a cancer mass and patient-derived cancer associated fibroblasts in a 3D vascularized colorectal cancer model. A total of six patient-derived CAF samples were successfully isolated and cultured from colorectal cancer tissue samples. Primary CAF characterisation is a topic of debate in literature. Some groups have done extensive characterisation on the gene and protein level of ‘CAF specific’ markers^18,51,29^. In this study, we successfully demonstrated that our CAF samples expressed widely recognised markers for CAF identification. CAF populations have often been classified based on their location within the tumour margin^52^. For comparison purposes, we used human dermal fibroblasts (HDFs) as our control or “healthy” stromal cells, which are not an immortalised cell line and could also start to differentiate into CAFs while in co-cultured with cancer cells within the tumouroids. It could be argued that most primary fibroblast samples will adapt a cancerous phenotype due to being cultured on plastic or in co-culture with cancer cells^53^. For future work it would be ideal to use paired CAF and NF samples from the same patient as part of a larger comparison study. Patient-derived NFs would serve as a better control and further our investigations into what signalling is caused by CAFs. Additionally, following on from this work potential of identifying what drives CAFs and their cancer promoting properties could be pin pointed. One pathway would be to generate knockout CAFs with a deletion of *HGF, TIMP1* and *FBLN5;* genes we found to be upregulated in the tumouroids and possibly responsible for increased invasion and vascular remodelling. Specific protagonists to VE-Cadherin such as could be used to potentially normalise the “leaky vasculature” caused by VEGFA upregulation, which are independent of one another.

## Acknowledgements

Judith Pape receives a stipend and EU fee funding from the EPSRC as part of the doctoral training program (DTP). Mark Emberton receives research support from the United Kingdom’s National Institute of Health Research (NIHR) UCLH/UCL Biomedical Research Centre and became an NIHR Senior Investigator in 2015. This work was funded by the NIHR Invention for Innovation (i4i) programme. Views expressed are those of the authors and not necessarily those of the NHS, the NIHR or the Department of Health.

I declare that the manuscript in its submitted form has been read and approved by all authors and is not being considered for publication elsewhere. The authors also declare no conflicts of interest.

